# Quenching corrinoid-based interactions in a model bacterial coculture

**DOI:** 10.64898/2025.12.31.697204

**Authors:** Zachary F. Hallberg, Zoila I. Alvarez-Aponte, Alison C. Gaudinier, Michiko E. Taga

## Abstract

Microbial community structure is driven, in part, by the metabolic interdependencies of resident microbes. Thus, manipulating specific metabolic interactions represents one attractive way to both understand how microbial communities perform complex functions and alter them for therapeutic or environmental effects. However, it is not yet possible to control the availability of those metabolites produced by some members of the community that are required by others. Here, we report the development of a metabolite ‘quenching’ strategy that disrupts a specific metabolic interaction involving corrinoids, the vitamin B_12_ family of cofactors, by applying a high-affinity corrinoid-binding protein, BtuG, to bacteria engaged corrinoid cross-feeding. Using a model coculture composed of *Sinorhizobium meliloti*, a bacterium that produces a corrinoid (cobalamin), and an *Escherichia coli* strain engineered to be corrinoid-dependent, we demonstrate corrinoid quenching by sequestration of extracellular corrinoid and show that BtuG specifically blocks corrinoid-dependent growth. We use this tool to calculate the amount of cobalamin released by *S. meliloti* cells and find that the cobalamin release rate is dependent on the growth phase of the producer, increasing to a maximum of approximately 40 cobalamin molecules per minute per cell in late exponential phase. This work establishes a strategy to selectively block microbial interactions that may be more broadly applied to dissecting community structure and function. We expect that applying high-affinity ‘molecular sponges’ to quench nutrient sharing will allow for the identification of key nutrients that structure microbial communities and empower precision microbiome manipulation strategies.

## Main Text

Nutrient competition and cross-feeding shape the function and composition of microbial communities (1, 2). Identifying nutrients that are key to structuring a given microbial community is critical to understanding how these ecological processes contribute to community composition. However, while altering the availability of an externally supplied nutrient such as a carbon source is straightforward, it remains difficult to control the availability of nutrients produced by and shared within a community. To address this limitation, we took inspiration from natural examples where metabolite availability is disrupted, or ‘quenched.’ In nutritional immunity, a host uses a high-affinity binding protein to sequester iron to limit pathogen growth (3, 4). Furthermore, some bacteria degrade quorum-sensing autoinducer signals to inhibit group behavior in competing microbes, a process termed quorum quenching (5). Developing methods to quench the availability of other metabolites, such as biosynthesized cofactors, would provide a powerful strategy to dissect nutrient interdependence and selectively control the growth of beneficial or pathogenic microbes.

Here, we describe a technique to quench interactions involving a family of biosynthesized micronutrients, corrinoids — the vitamin B_12_ (cobalamin) family of enzyme cofactors. Genomic data predicts that corrinoids are synthesized by only 36% of bacterial species, but used by over 80% (6-9), suggesting that they are important shared metabolites in microbial communities. We and others have shown that corrinoid addition to microbial communities influences community structure (10-13), demonstrating that corrinoids have the ability to alter the composition and growth of microbial communities. Nevertheless, direct demonstration of their importance as naturally-shared metabolites in a community context remains lacking, and could be performed with a suitable corrinoid quenching method. To demonstrate that corrinoid interactions can be quenched, we used BtuG, a high-affinity corrinoid-binding protein required for corrinoid uptake in *Bacteroides thetaiotaomicron* (14, 15). This quenching method offers a proof-of-concept that can be applied to manipulate interactions involving other shared nutrients.

We first established a model coculture consisting of corrinoid-producing and -dependent bacteria. As the producer, we selected *Sinorhizobium meliloti* Rm1021 (*Sm*), because unlike some other bacteria, it produces a corrinoid (cobalamin) at levels that support dependent growth, and some of this cobalamin is released into culture supernatants (**Figure S1A**) for potential use by corrinoid-dependent microbes in coculture (**Figure S1B**). For the corrinoid-dependent strain, we used a corrinoid-dependent *E. coli* mutant expressing a fluorescent protein for growth readout (*ΔmetE gfp*^*+*^, abbreviated as *Ec*). This strain has a deletion of *metE*, making it reliant on a corrinoid for methionine synthesis when no methionine is supplied. To establish the growth response of the two strains in mono- and coculture, we tracked growth by measuring O.D._600_ and fluorescence (**Figure 1**). We screened multiple carbon sources for the ability to support growth of the two strains. Some carbon sources supported monoculture growth of both species (Groups 1-2), while others supported only one (Groups 3-4). Coculture growth was observed in a subset of carbon sources that support growth of both taxa (Group 1), and in one carbon source (lactose) that only the dependent could metabolize (Group 4). We selected glycerol as the carbon source for subsequent experiments, as neither strain exhibits a distinct growth advantage in this medium, and *E. coli* does not undergo the acetate switch when grown in this carbon source (16).

**Figure 1.**
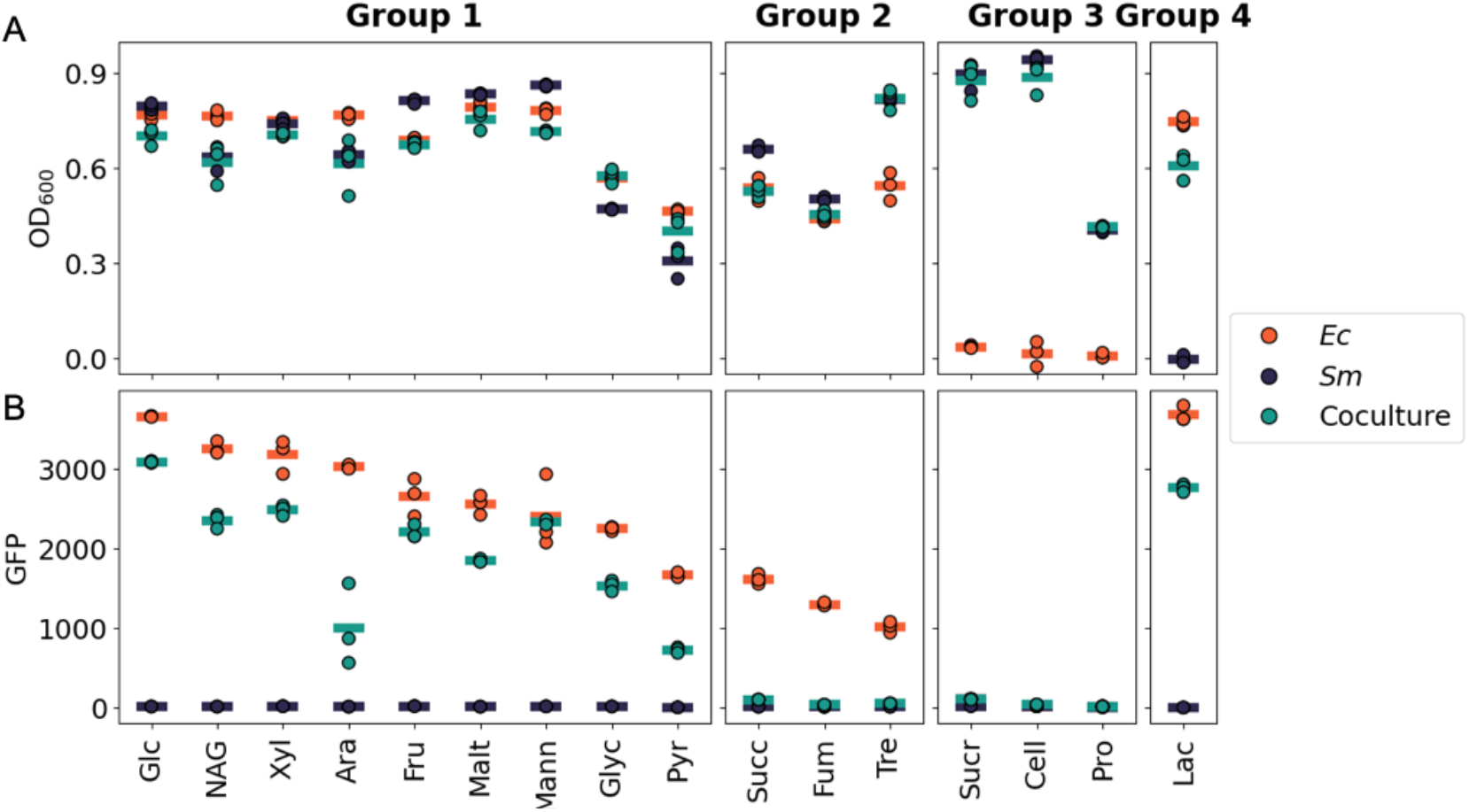
Growth of *Sm* (producer) and *Ec* (dependent) in mono- and coculture across carbon sources. (A) O.D._600_ and (B) GFP measured at 48 h are shown for *Sm* in monoculture, GFP-labeled *Ec* supplemented with cyanocobalamin (vitamin B_12_), and cocultures of both grown in M9 medium with the labeled additive as the carbon source. Carbon sources are separated by culture response: Group 1 - carbon sources in which both strains grow that elicit dependent growth in coculture, Group 2 - carbon sources in which both strains grow but elicit no dependent growth in coculture. Groups 3 and 4 represent carbon sources in which only *Sm* (Group 3) or *Ec* can grow in monoculture (Group 4). The values shown are O.D._600_ and GFP measurements with uninoculated controls subtracted. Average values for triplicate biological samples are shown as a horizontal line. Carbon sources are abbreviated as: Glc = Glucose, NAG = N-acetylglucosamine, Xyl = Xylose, Ara = Arabinose, Fru = Fructose, Malt = Maltose, Mann = Mannitol, Glyc = Glycerol, Pyr = Pyruvate, Succ = Succinate, Fum = Fumarate, Tre = Trehalose, Sucr = Sucrose, Cell = Cellobiose, Pro = Proline, Lac = Lactose.

A successful corrinoid-quenching strategy must specifically prevent corrinoid uptake without interfering with other physiological processes. To test specificity of BtuG as a corrinoid quencher, we first determined whether the addition of purified BtuG protein to *Ec* grown in monoculture could specifically inhibit corrinoid-dependent growth. When cultured in 1 nM cobalamin with increasing concentrations of purified BtuG, we found that growth was inhibited at concentrations higher than 1.23 nM (**Figure S1C**). This BtuG-dependent growth suppression is specific to cobalamin metabolism, because when methionine is supplied instead of cobalamin, growth remains robust at high BtuG concentrations (**Figure S1C**).

Having determined that BtuG can quench cobalamin-dependent growth in monoculture when cobalamin is supplied in the growth medium, we tested whether BtuG could affect *Ec* growth when cobalamin is released by *Sm*. Indeed, we observed that, when cocultured with *Sm* in media without added cobalamin, *Ec* growth was suppressed by addition of BtuG, with higher concentrations completely inhibiting the growth of *Ec* (**Figure 2A, S2A**). This result provides a proof-of-concept that BtuG can be used as a tool to block corrinoid sharing interactions.

**Figure 2.**
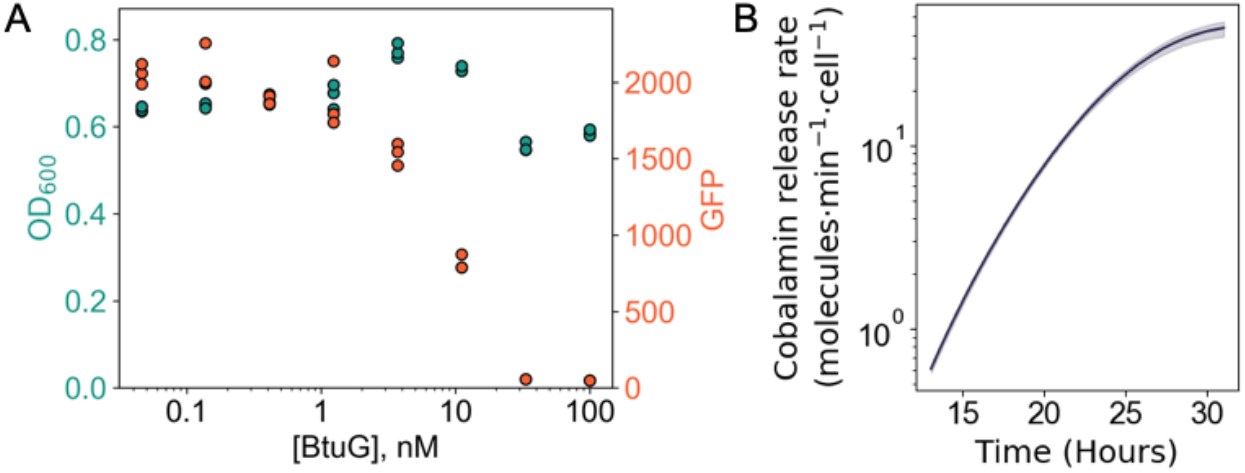
Coculture response to corrinoid quenching. (A) O.D._600_ and GFP signal after 48 h from cocultures of *Sm* and *Ec* with increasing concentrations of BtuG. (B) Modeled change in cobalamin-providing rate by *Sm* per cell obtained using the change in lag phase in higher BtuG concentrations. Monoculture growth of *Sm* was approximated by the O.D._600_ at 100 nM BtuG, where *Ec* growth was inhibited.

In addition to providing evidence that corrinoids are released by *Sm* prior to uptake by *Ec*, our quenching data provided a novel way to estimate the corrinoid release rate by *Sm*. At different BtuG concentrations, *Ec* exhibited different lag phase durations (**Figure S2B**). We reasoned that this lag phase represents the time required for *Sm* to produce and release enough cobalamin to saturate the added BtuG and support cobalamin-dependent growth of *Ec*. By correlating the concentration of added BtuG (corresponding to the amount of cobalamin sequestered) with the *Ec* lag phase duration, we calculated the amount of cobalamin released by *Sm* over time (**Figure 2B**). Based on these calculations, we find the cobalamin release rate during *Sm* lag phase is low (<1 molecule·min^−1^·cell^−1^). As *Sm* progresses through exponential phase, cobalamin release increases, reaching almost 40 molecules·min^-1^·cell^-1^. Thus, we conclude that cobalamin release by *Sm* is, in part, a function of growth phase.

Here, we have developed a strategy to sequester corrinoids using a high-affinity corrinoid-binding protein, demonstrating an extracellular corrinoid ‘sharing’ interaction between *Sm* and *E*c in coculture. We also showed that *Sm* has a higher rate of corrinoid release during late exponential growth. Our finding that *B. thetaiotaomicron* BtuG can quench a corrinoid-sharing interaction suggests that *Bacteroides* may use BtuG both for corrinoid uptake and to engage in corrinoid quenching as a mechanism to compete with other microbes in the gut. The nutrient quenching technique developed here offers a strategy to investigate metabolite cross-feeding interactions more broadly. Other high-affinity metabolite-binding proteins could be used as a general tool to investigate the roles of diverse nutrients in shaping microbial community structure, further enabling precision modulation of microbial communities.

## Data Availability

All raw data and associated code is available in the supplementary files.

## Acknowledgements and Funding

We thank Andy Goodman for providing the BtuG expression plasmid used in this work. We thank members of the Taga lab for their constructive comments. This work was supported by the U.S. Department of Energy (DOE), Office of Science, Office of Biological and Environmental Research, Genomic Science Program under Award Number DE-SC0020155 to M.E.T., National Institutes of Health (NIH) award R35GM139633 to M.E.T., and NIH award K99GM143653 to Z.F.H.

## Notes

### Competing Interest Statement

The authors have declared no competing interest.

