## Supplementary Data and Figures for "Quenching corrinoid-based interactions in a model bacterial coculture"

### Materials and Methods.

#### Strain construction.

*E. coli* strain MG1655 *ΔmetE ΔnfsA::RFP-GFP-kan$^R$* was constructed by P1 transduction of the *ΔnfsA::RFP-GFP-kan$^R$* from MG1655 *ΔnfsA::RFP-GFP-kan$^R$* (1) into *E. coli* strain MG1655 *ΔmetE*.

#### BtuG expression and purification.

A signal sequence truncation mutant of BtuG2 from *Bacteroides thetaiotaomicron* VPI-5482 with a C-terminal $His_{10}$-tag in a pET21-derived plasmid was expressed and purified as previously reported (2, 3). Protein aliquots were diluted in storage buffer (1×PBS, pH 8.0, composed of 10.1 mM $Na_2HPO_4$, 1.8 mM $KH_2PO_4$, 137 mM NaCl, 2.7 mM KCl) to a 10x working concentration for experiments.

#### Strain preparation and preculturing.

For preparation of cells for all experiments, strains were streaked from frozen (-80$^o$C) 25% glycerol stocks (*E. coli*, *P. denitrificans*) or 10% DMSO stocks (*S. meliloti*) onto LB-agar (1.5%) plates to obtain single colonies. Individual colonies were inoculated into M9-glucose (0.2%) medium supplemented with 1× SL10 minerals, 1× Wolin's vitamins (without $B_{12}$), and, for *E. coli* Δ*metE* strains, methionine (1 mg/mL) (4, 5). Cultures were incubated at 30°C for 48 h with shaking at 200 revolutions per minute (rpm). Cells were pelleted by centrifugation (5000 × rcf, 4 min) of 1 mL of saturated culture, the supernatant aspirated, and resuspended in 1× PBS (1 mL, pH 7.4). Cells were washed a total of three times in this way.

#### Microbial Growth Assays.

All growth assays were conducted in 96-well microtiter plates (Corning Costar Assay Plate 3904) and sealed with air-permeable membranes (Breathe-Easy, Diversified Biotech). Cultures were incubated at 30°C for 24-48 h, either in a bench top heated plate shaker (Southwest Science) at 1,200 rpm, or a Biotek Synergy 2 plate reader with linear shaking at 1,140 cycles per minute. $O.D._{600}$ values and GFP fluorescence (excitation: 485/20, emission: 528/20) were measured in the Biotek Synergy 2 plate reader. Data were plotted and analyzed in Python.

For initial coculture screens, 2 µL of prepared producer culture and 2 µL of the dependent strain (if applicable) were added to 200 µL of modified M9 medium as above or supplemented with 1× of the 19 amino acids used for EZ-rich medium (6). For coculture experiments screening carbon sources, cells were resuspended and combined in M9 medium lacking a carbon source so that both strains were at a starting $O.D._{600}$ of 0.001. 180 µL of this cell mixture was added to 20 µL of carbon source (2% w/v, for a 0.2% final concentration). For controls, 20 µL of 1× PBS was used as a negative control, and 20 µL of 10 nM cyanocobalamin was added for the $B_{12}$ control (final concentration 1 nM). To determine BtuG quenching efficacy, 2 µL of washed *E. coli* culture (diluted to an $O.D._{600}$ of 0.1 in 1× PBS) were added to 180 µL of modified M9-glycerol (0.4%) medium (supplemented with either methionine (1 mg/mL) or vitamin $B_{12}$ (1 nM)) and 20 µL of BtuG at a 10× concentration. For BtuG coculture experiments, 2 µL of prepared *S. meliloti* cells and 2 µL of prepared *E. coli* cells (each diluted to an $O.D._{600}$ of 0.1 in 1× PBS) were added to 180 µL of modified M9-glycerol (0.4%) and 20 µL of BtuG at a 10× stock concentration.

#### Calculating rate of cobalamin release per cell.

To calculate the amount of cobalamin released per cell per time, growth curves were analyzed in Python using a custom script to extract lag phase (as analyzed by GFP signal for *E. coli* bioassay growth), and were confirmed using AMiGA (7). The amount of cobalamin released over the *E. coli* lag periods was calculated by dividing the molar amount of BtuG added by the difference in lag phase duration observed at a given BtuG concentration relative to the lag phase with no BtuG added. These values were fit to an exponential curve, and the derivative of this curve was taken to obtain the instantaneous rate of cobalamin release per unit time. This value was then divided by the number of *S. meliloti* cells at that same timepoint (obtained from the $O.D._{600}$ of the producer-only growth curve, i.e., the 100 nM BtuG condition, and assuming an $O.D._{600} = 1$ corresponds to $10^9$ cells/mL) to obtain the rate of cobalamin released per cell per time.

**Supplemental Table 1.** List of bacterial strains used in this study.

| Species | Strain | Genotype | Reference |
|---|---|---|---|
| *E. coli* | MG1655 *ΔmetE* | *ΔmetE::FRT* | (Mok et al, 2022) |
| *E. coli* | MG1655 *ΔmetE nfsA::RFP-GFP-kan* | *ΔmetE::FRT nfsA::*$P_{J23119}$*-RFP-*$P_{J23119}$*-GFP-kan* | This work |
| *E. coli* | MG1655 *nfsA::RFP-GFP-kan* | *nfsA::*$P_{J23119}$*-RFP-*$P_{J23119}$*-GFP-kan* | (Qi et al, 2013) |
| *S. meliloti* | Rm1021 | | (Meade et al, 1982) |
| *P. denitrificans* | ATCC 13867 | | (Ainala et al, 2013) |

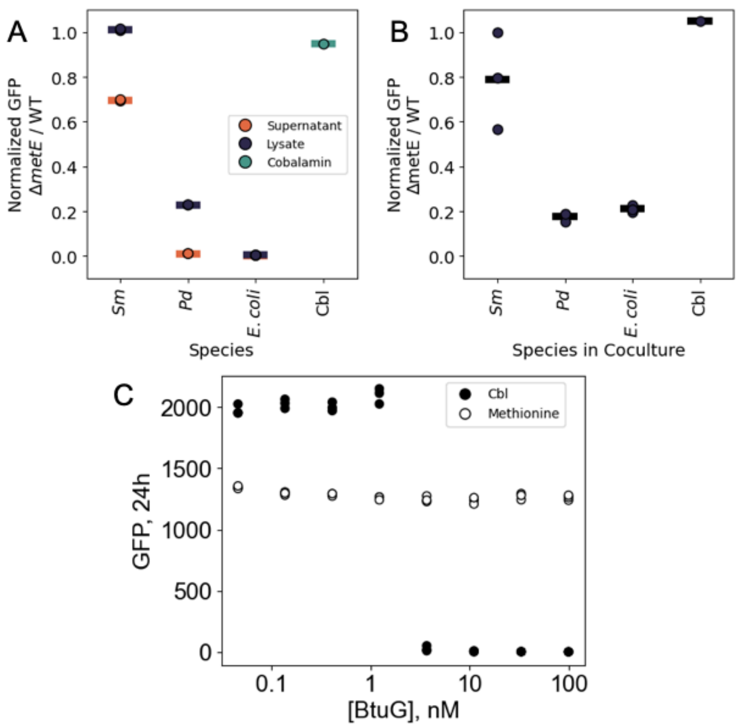


**Figure S1. Response of *Ec* to *Sm* and BtuG.** (A) Growth of *E. coli ΔmetE gfp*+ reporter strain (*Ec*), measured by GFP fluorescence in the presence of supernatant or cell lysate of corrinoid producers *S. meliloti* (*Sm*) and *Pseudomonas denitrificans* (*Pd*), and non-producer *E. coli* wild type strain MG1655 or 1 nM cyanocobalamin (Cbl). Values are normalized to the GFP signal from *Ec* in the same conditions (2 biological replicates), and horizontal lines refer to average values of all replicates. (B) Fluorescence of cocultures containing the indicated bacteria with *Ec* (3 biological replicates). Horizontal lines refer to average values of all replicates. (C) GFP signal of *Ec* supplemented with methionine (1 mg/mL) or cyanocobalamin (1 nM) and increasing concentrations of purified BtuG. We note that GFP fluorescence is consistently lower when cultures are supplemented with methionine than with cobalamin (3 biological replicates).

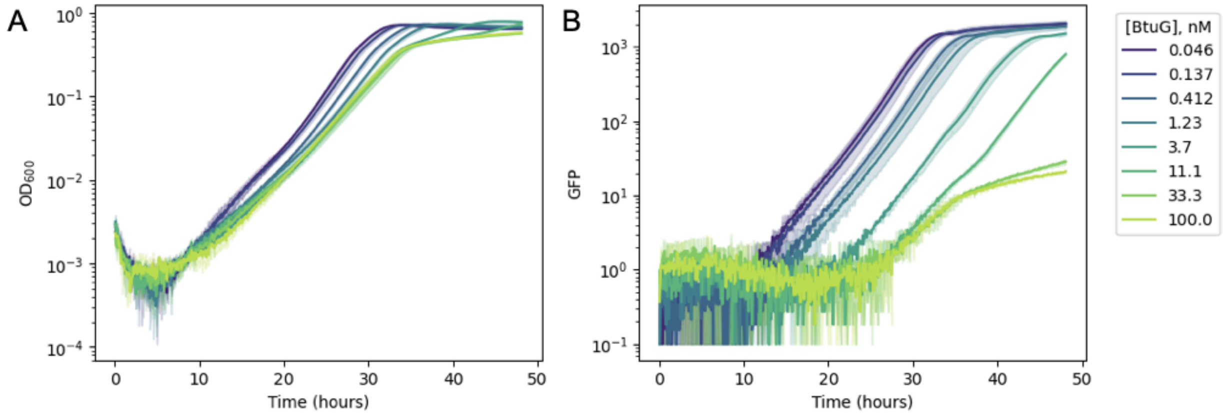


**Figure S2. Dynamic response of cobalamin cross-feeding cocultures to BtuG.** O.D.$_{600}$ (A) and GFP fluorescence (B) of cocultures containing *Sm* and *Ec* supplemented with increasing concentrations of corrinoid-quenching protein BtuG over time). Measurements of uninoculated samples were background subtracted and normalized to and initial O.D.$_{600}$ of 0.001 (A) and an initial GFP signal of 1 (B).
